# Distance from Sub-Saharan Africa Predicts Mutational Load in Diverse Human Genomes

**DOI:** 10.1101/019711

**Authors:** Brenna M. Henn, Laura R. Botigué, Stephan Peischl, Isabelle Dupanloup, Mikhail Lipatov, Brian K. Maples, Alicia R. Martin, Shaila Musharoff, Howard Cann, Michael Snyder, Laurent Excoffier, Jeffrey M. Kidd, Carlos D. Bustamante

## Abstract

The Out-of-Africa (OOA) dispersal ~50,000 years ago is characterized by a series of founder events as modern humans expanded into multiple continents. Population genetics theory predicts an increase of mutational load in populations undergoing serial founder effects during range expansions. To test this hypothesis, we have sequenced full genomes and high-coverage exomes from 7 geographically divergent human populations from Namibia, Congo, Algeria, Pakistan, Cambodia, Siberia and Mexico. We find that individual genomes vary modestly in the overall number of predicted deleterious alleles. We show via spatially explicit simulations that the observed distribution of deleterious allele frequencies is consistent with the OOA dispersal, particularly under a model where deleterious mutations are recessive. We conclude that there is a strong signal of purifying selection at conserved genomic positions within Africa, but that many predicted deleterious mutations have evolved as if they were neutral during the expansion out of Africa. Under a model where selection is inversely related to dominance, we show that OOA populations are likely to have a higher mutation load due to increased allele frequencies of nearly neutral variants that are recessive or partially recessive.

## Significance statement

Human genomes carry hundreds of mutations that are predicted to be deleterious in some environments, potentially impacting the health or fitness of an individual We characterize the distribution of deleterious mutations among diverse human populations, modeled under different selection coefficients and dominance parameters. Using a new dataset of diverse human genomes from seven different populations, we use spatially explicit simulations to reveal that classes of deleterious alleles have very different patterns across populations reflecting the interaction between genetic drift and purifying selection. We show that there is a strong signal of purifying selection at conserved genomic positions within African populations, but most predicted deleterious mutations have evolved as if they were neutral during the expansion out of Africa.

## Introduction

It has long been recognized that a human genome may carry many strongly deleterious mutations; Morton et al (1) estimated that each human carries on average 4-5 mutations that would have a “conspicuous effect on fitness” if expressed in a homozygous state. Empirically estimating the deleterious mutation burden is now feasible through next-generation sequencing (NGS) technology, which can assay the complete breadth of variants in a human genome. For example, recent sequencing of over 6,000 exomes revealed that nearly half of all surveyed individuals carried a likely pathogenic allele in a known Mendelian disease gene (i.e. from a disease panel used for newborn screening) (2). While there is some variation across individuals in the number of deleterious alleles per genome, we still do not know if there are significant differences in deleterious variation among populations. Human populations vary dramatically in their levels of neutral genetic diversity, which suggests variation in the effective population size, *N_e_.* Theory suggests that the efficacy of natural selection is reduced in populations with lower *N_e_* because they experience greater genetic drift (3, 4). In an idealized population of constant size, the efficacy of purifying selection depends on the relationship between *N_e_* and the selection coefficient ‘*s*’ against deleterious mutations. If *4N_e_s* << 1, deleterious alleles evolve as if they were neutral and can, thus, reach appreciable frequencies. This theory raises the question of whether human populations carry differential burdens of deleterious alleles due to differences in demographic history.

Several recent papers have tested for differences in the burden of deleterious alleles among populations; these papers have focused on primarily comparing populations of western European and western African ancestry. Despite similar genomic datasets, these papers have reached a variety of contradictory conclusions (4–9). Initially, Lohmueller et al. (10) found that a panel of European-Americans carried proportionally more derived, deleterious alleles than a panel of African-Americans, potentially as the result of the Out-of-Africa bottleneck. More recently, analyses using NGS exome datasets from samples of analogous continental ancestry found small or no differences in the average number of deleterious alleles per genome between African-Americans and European-Americans – depending on which prediction algorithm was used (11–13). Simulations by Fu et al. (11) found strong bottlenecks with recovery could recapitulate patterns of differences in the number of deleterious alleles between African and non-African populations, supporting Lohmueller et al. (10), but in contrast to work by Simons et al. (12).

It is important to note two facts about these contradictory observations: first, these papers tend to use different statistics, which differ in power to detect changes across populations, as well as the impact of recent demographic history (6, 11). Lohmueller et al. (10) compared the relative number of non-synonymous to synonymous (or “probably damaging” to “benign”) SNPs per population in a sample of *n* chromosomes whereas Simons et al. (12) examined the special case of *n* = 2 chromosomes; namely, the average number of predicted deleterious alleles per genome (i.e., heterozygous + 2 * homozygous derived variants per genome). One way to think about these statistics is that the total number of variants, S, gives equal weight, *w* = 1, to a SNP regardless of its frequency, *p*. The average number of deleterious variants statistic gives weights proportional to the expected heterozygous and homozygous frequencies or *w* = 2*p*(1-*p*) + *p*^2^ = 2*p* − *p*^2^. The average number of deleterious alleles per genome is fairly insensitive to differences in demographic history since heterozygosity is biased towards common variants. In contrast, the proportion of deleterious alleles has greater power to detect the impact of recent demographic history for large *n* across the populations since it is sensitive to rare variants which tend to be more numerous, younger, and enriched for functionally important mutations (14–16). Secondly, empirical comparisons between two populations have focused primarily on an additive model for deleterious mutations, even though there is evidence for pathogenic mutations exhibiting a recessive or dominant effect (17, 18), and possibly an inverse relationship between the strength of selection *s* and the dominance parameter *h* (19).

There remains substantial conceptual and empirical uncertainty surrounding the processes that shape the distribution of deleterious variation across human populations. We aim here to clarify three aspects underlying this controversy: 1) are there empirical differences in the total number of deleterious alleles among multiple human populations?, 2) which model of dominance is appropriate for deleterious alleles (i.e., should zygosity be considered in load calculations)?, 3) are the observed patterns consistent with predictions from models of range expansions accompanied by founder effects? We address these questions with a new genomic dataset of seven globally distributed human populations.

## Results

*Population history and global patterns of genetic diversity:* We obtained moderate coverage whole-genome sequence (median depth 7x) and high coverage exome sequence data (median depth 78x) from individuals from seven populations from the Human Genome Diversity Panel (HGDP) (20) Unrelated individuals (no relationship closer than 1^st^ cousin) were selected from seven populations chosen to represent the spectrum of human genetic variation from throughout Africa and the Out-of-Africa expansion, including individuals from the Namibian San, Mbuti Pygmy (Democratic Republic of Congo), Algerian Mozabite, Pakistani Pathan, Cambodian, Siberian Yakut and Mexican Mayan populations (**Fig. 1A, Table S1**). The 2.48 Gb full genome callset consisted of 14,776,723 single nucleotide autosomal variants, for which we could orient 97% to ancestral/derived allele status (*Supplementary Note*).

**Figure 1:**
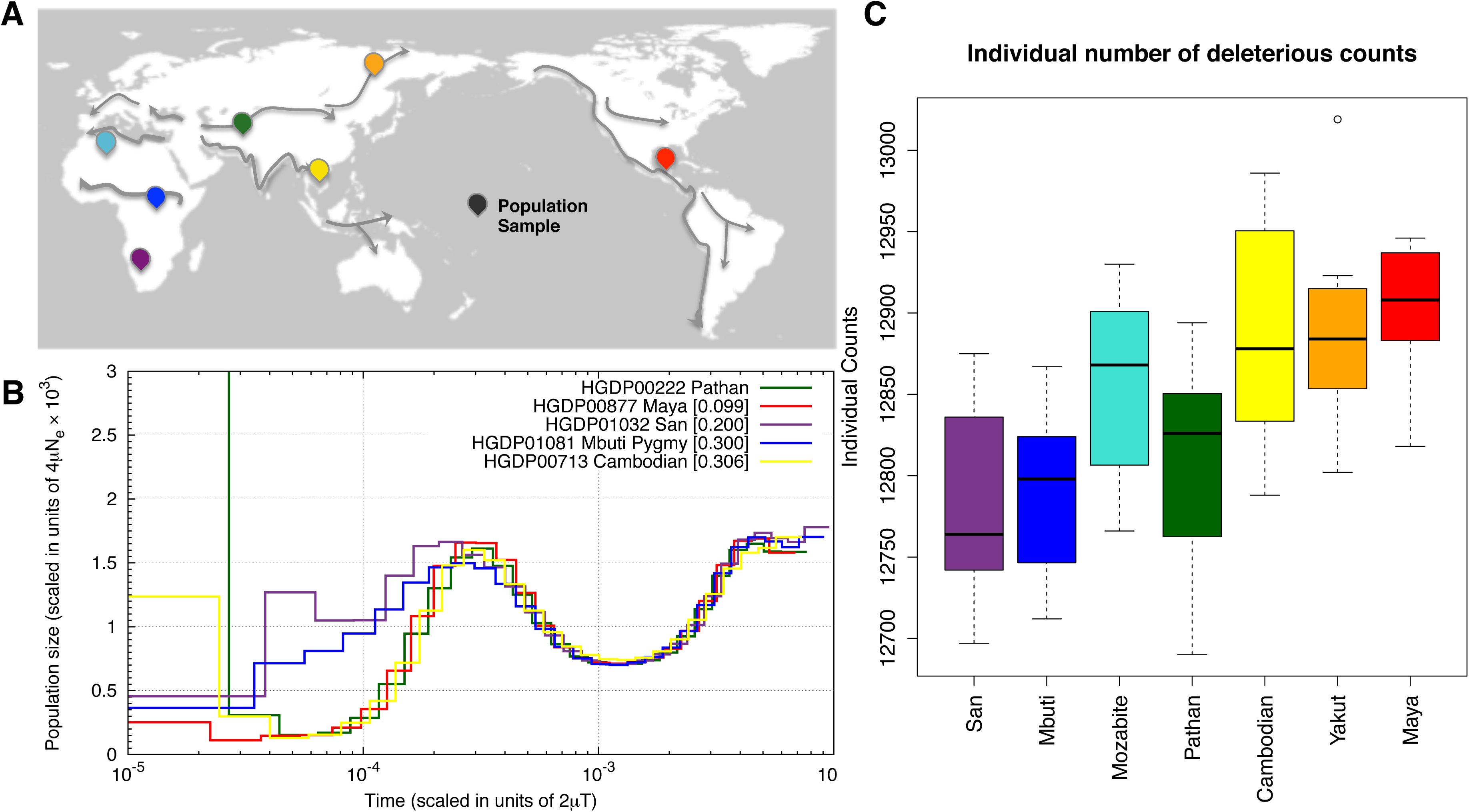
Decrease in heterozygosity and estimated N_e_ with distance from southern Africa. **A**) Locations of HGDP populations sampled for genome and exome sequencing are indicated on the map. Putative migration paths after the origin of modern humans are indicated with arrows (adapted from(6)). **B**) PSMC curves for individual genomes, corrected for differences in coverage. While populations experiencing an Out of Africa bottleneck have substantially reduced *N_e_,* African populations also display a reduction in *N_e_* between ~100*kya* and 30*kya,* (see **Supplementary Results** for simulations of population history with resulting PSMC curves.) **C**) For each individual’s exome, the number of putatively deleterious variants (equivalent to number of heterozygotes + twice the number of derived homozygotes) is shown by population.

Heterozygosity among the 7 populations decreases with distance from southern Africa, consistent with an expansion of humans from that region (21). The Namibian San population carried the highest number of derived heterozygotes ~2.39 million per sample, followed closely by the Mbuti Pygmies (**Table S1, Fig. S5**). The North African Mozabites carry more heterozygotes than the Out-of-Africa populations in our dataset (2 million), but substantially fewer than the sub-Saharan samples, likely reflecting a complex history of an Out-of-Africa migration, followed by re-entry into North Africa and subsequent recent gene flow with neighboring African populations (22). The Maya have the lowest median number of heterozygotes in our sample, ~1.5 million, which may be inflated due to recent European admixture (23). Two Mayan individuals displayed substantial recent European admixture (>20%) as assessed with local ancestry assignment (24) (**Fig. S6**); these individuals were removed from analyses of deleterious variants. When we recalculated heterozygosity in the Maya, it was reduced by 3.5%. The decline in heterozygosity in OOA populations with distance from Africa strongly supports earlier results based on SNP array and microsatellite data for a serial founder effect model for the OOA dispersal (25, 26). We analyzed population history for individuals having sufficient coverage from five of the studied populations using the pairwise sequential Markovian coalescent software (PSMC) to estimate changes in *N_e_* (11, 12, 27). Since dating demographic events with PSMC is dependent on both the assumed mutation rate and the precision with which a given event can be inferred, we compare relative bottleneck magnitudes and timing among the seven HGDP populations. Consistent with previous analyses (27), the Out-of-Africa populations show a sharp reduction in *N_e_,* with virtually identical population histories (**Fig. 1B,** *Supplementary Note*). Simulations indicate that the magnitude of the twelvefold bottleneck is accurately estimated (**Fig. S7**), even if the time of the presumed bottleneck is difficult to estimate precisely using PSMC. Interestingly, both the Mbuti and the Namibian San show a moderate reduction in *N_e_* relative to the ancestral maximum, with the San experiencing an almost two-fold reduction in *N_e_* and the Mbuti display a reduction intermediate between the San and OOA populations (see also (20, 28, 29)). These patterns are consistent with multiple population histories (e.g. both short and long bottlenecks) and multiple demographic events, including a reduction in sub-structure from the ancestral human population rather than a bottleneck *per se* (27).

*Differences in Deleterious Alleles per Individual Genomes:* Due to differences in coverage among the whole genome sequences, our subsequent analyses focus on the high-coverage exome dataset (78x median coverage) in order to minimize any bias in comparing populations (**Table S1**, *Methods*). We classified all mutations discovered in the exome dataset into categories based on GERP Rejected Substitution (RS) scores. These conservation scores reflect various levels of constraint within a mammalian phylogeny (*Methods*) and are used to categorize mutations by their predicted deleterious effect (30, 31). Importantly, the allele present in the human reference genome was not used in the GERP RS calculation, avoiding the reference-bias effect previously observed in other algorithms (11, 12) (**Fig. S8A**). Variants were sorted into four groups reflecting the likely severity of mutational effects: “neutral” (-2 < GERP < 2), “moderate” (2 ≤ GERP ≤ 4), “large” (4 < GERP < 6) and “extreme” (GERP ≥ 6) *(Supplementary Note,* **Fig. S9**). GERP categories were concordant with ANNOVAR functional annotations (**Table S2, Fig. S8B**).

When considering the total number of derived alleles per individual, defined here as: A_I_ = (1 × HET) + (2 × HOM_der_), we observe an increase of predicted deleterious alleles with distance from Africa (**Fig. 1C**). The number of predicted deleterious alleles per individual increases along the range expansion axis (from San to Maya), consistent with theoretical predictions for expansion load (32). The maximal difference in the number of deleterious alleles between African and OOA individuals is ~150 alleles. This result is consistent with theoretical predictions; the rate at which deleterious mutations accumulate in wave front populations is limited by the total number of mutations occurring during the expansion (32). Assuming an exomic mutation rate of *u* = 0.5 per haploid exome and an expansion that lasted for *t* = 1,000 generations, a very conservative upper limit for the excess of deleterious alleles in OOA individuals would be 2\**u*t* = 1000. The cline in A_I_ is most pronounced for large effect alleles (4 < GERP < 6, **Fig. 2E**), whereby the San individuals carry A_I_ = 4,450 large effect alleles on average, increasing gradually to 4,550 in Yakut. The Mayans carry slightly fewer large effect mutations per individual than the Yakut, which may be influenced by the residual European ancestry (between 5%-20%) in our sample. For extreme alleles (GERP ≥ 6), each individual in the dataset carries on average 110–120 predicted highly deleterious alleles with no significant differences among populations (**Fig. 2F**). The average additive GERP score - obtained by counting the GERP scores at homozygous sites twice - for all predicted deleterious variants per individual is lowest in the San (~3.3) and highest in the Maya (~3.8).

**Figure 2:**
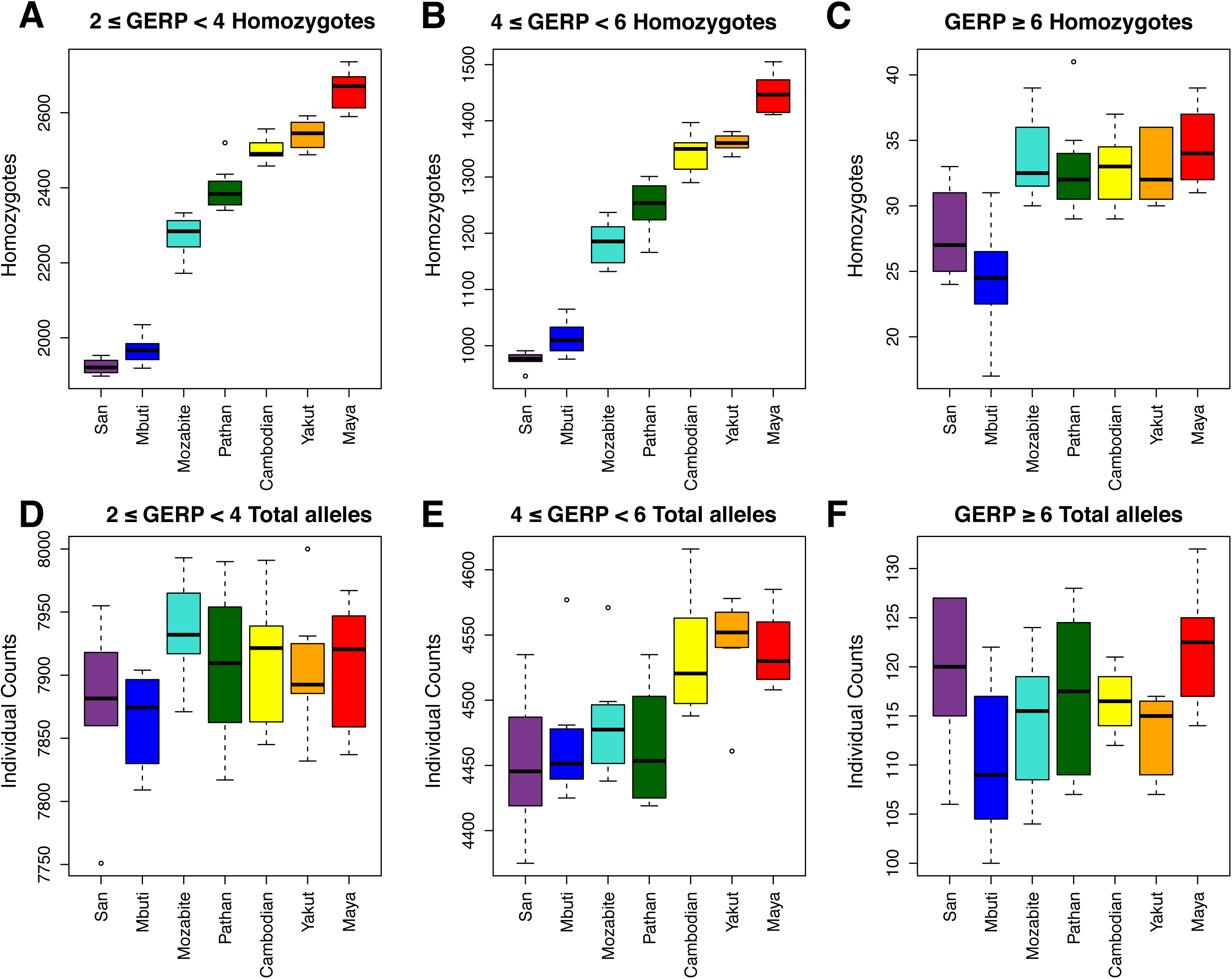
Individual counts of deleterious variants. (**A-C**) For each individual’s exome, the number of derived homozygotes is plotted by population for moderate, large and extreme effect GERP categories. (**D-F**) For each individual’s exome, the number of derived variants (equivalent to number of heterozygotes + twice the number of homozygotes) is plotted by population for moderate, large and extreme effect GERP categories.

Similar patterns are found when we consider the number of derived homozygous sites per individual. We find that individuals from OOA populations exhibit significantly more homozygotes for moderate, large and extreme variants than African populations (**Fig. 2 A-C**). In addition, we observe a clear increase in the number of derived homozygotes with distance from Africa for moderate (2 ≤ GERP ≤ 4) and large (4 < GERP < 6) mutation effects categories, whereas the number of derived “extreme” homozygotes (GERP ≥ 6) is similar among OOA populations: all OOA genomes possess 30–40 extremely deleterious alleles in homozygous state (**Fig. 2C**). These patterns are in excellent agreement with theoretical predictions for the evolution of genetic variation during range expansions (7). The average GERP score per individual for derived homozygous variants is less differentiated than the additive model (above), varying between 2.43 to 2.49.

It is important to note that A_I_ is strongly influenced by common variants. Goode et al. (33) observed that as much as 90% of deleterious alleles in a single genome have a derived allele frequency greater than 5%, suggesting that the bulk of mutational burden using this metric will come from common variants. To explore this idea, we randomly chose an individual in each population and calculated the proportion of deleterious variants that are rare (<10%, i.e. a singleton within our population samples) and common (>10%), for each GERP category (**Fig. 3A**). Common deleterious alleles contribute to more than 90% of an individual’s A_I_, and the proportion of common deleterious variants increases with distance from Africa, as can be seen by the decrease of rare deleterious variants. This includes common large effect variants, which make up proportionally more of A_I_ for an OOA individual than for an African individual: for example, in a Mayan individual, 93% of large effect variants are common compared to a San individual, where only 85% of large effect variants are common (**Fig. S12**). Given the small number of chromosomes in each population (*n*=14−16), estimates of allele frequencies are subject to sampling effects. We recently performed the same analysis on exome data from the 1000 Genome Phase 1 Project (34). We find a similar pattern as in our HGDP data: on a per genome basis, common variants represent a majority of the alleles predicted to be deleterious (5).

**Figure 3:**
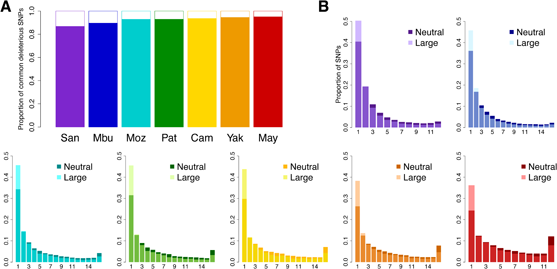
Differences in the proportion of deleterious alleles by frequency class. A) The proportion of rare versus common deleterious variants per individual. For a given individual, deleterious variants were divided into common (>10%, solid colors) and rare (<10%, white space). The contribution of common deleterious variants to an individual’s burden is much greater than rare variants. B) For each population, we calculated the proportional site frequency spectrum by plotting the proportion of deleterious large effect alleles in each frequency class (translucent coloring) along with the proportion of neutral alleles for each frequency class (opaque coloring). African populations have proportionally fewer rare deleterious alleles than expected from neutrality. Populations with OOA ancestry have proportionally more fixed deleterious mutations.

*Differences in deleterious alleles at the population level:* To further elucidate the relationship between predicted mutation effect and allele frequencies, we compared the site frequency spectrum (SFS) for neutral and large (4 < GERP < 6) effect variants (**Fig. 3B, see Fig. S14** for a comparison between neutral and extreme variants). For all populations, singletons are enriched for deleterious variants (as compared to neutral variants), consistent with the effect of purifying selection against deleterious variants (15, 35). However, the SFSs of OOA and African populations show marked differences. The neutral and deleterious SFS of OOA populations show a global shift towards higher frequencies, consistent with the effects of serial bottlenecks/founder effects. It follows that OOA populations have fewer rare deleterious variants than Africans, as well as a larger proportion of fixed deleterious alleles; almost 7.9% of large effect variants are fixed in the Maya, whereas the San have only 1.8% of deleterious variants fixed (**Fig. 3B**).

*Simulations of purifying selection under a range expansion:* We sought to interpret the population-specific patterns of genetic diversity for each GERP category under a model including serial founder effects across geographic space and purifying selection. We simulated the evolution of both neutral and deleterious mutations under a simple model of range expansion in a two dimensional habitat (see **Fig. S21**). At selected loci, the ancestral allele was assumed selectively neutral and mutants reduced an individual’s fitness by a factor 1-*s* only if it was present in homozygous state, that is, deleterious mutations were assumed to be completely recessive. Three thousand generations (corresponding to about 75 *kya*) after the onset of the range expansion, we computed the average expected heterozygosity for all populations. Computational limitations of individual-based simulations prohibit a complete exploration of the parameter space for this model but, by varying migration rates and selection coefficients, we identified parameter values that fit the observed clines in heterozygosity reasonably well (**Fig. 4B**). Specifically, we first identified selection coefficients that yield the same relative differences between observed neutral and selected heterozygosities (**Fig. 4A**). Then, the migration rate was adjusted to fit the observed clines in heterozygosities, assuming that the distance between two demes is 250 km (**Fig. 4B**). The fit selection coefficients were 0, 1.25×10^−4^, 1×10^−3^ and 2×10^−3^ for neutral, moderate, large, and extreme GERP scores categories, respectively; the GERP ≥ 6 category showed the worst fit and observed counts indicate that even stronger selection coefficients should be considered for these extreme mutations (16). We performed the same analysis using a model in which mutations are co-dominant and, as expected, we found that the fit selection coefficients are smaller than those obtained a recessive model. These coefficients are estimated as: *s* = 0, 0.5×10^−4^, 1.2×10^−4^ and 2×10^−4^, respectively (see **Fig. S16**) (16).

**Figure 4:**
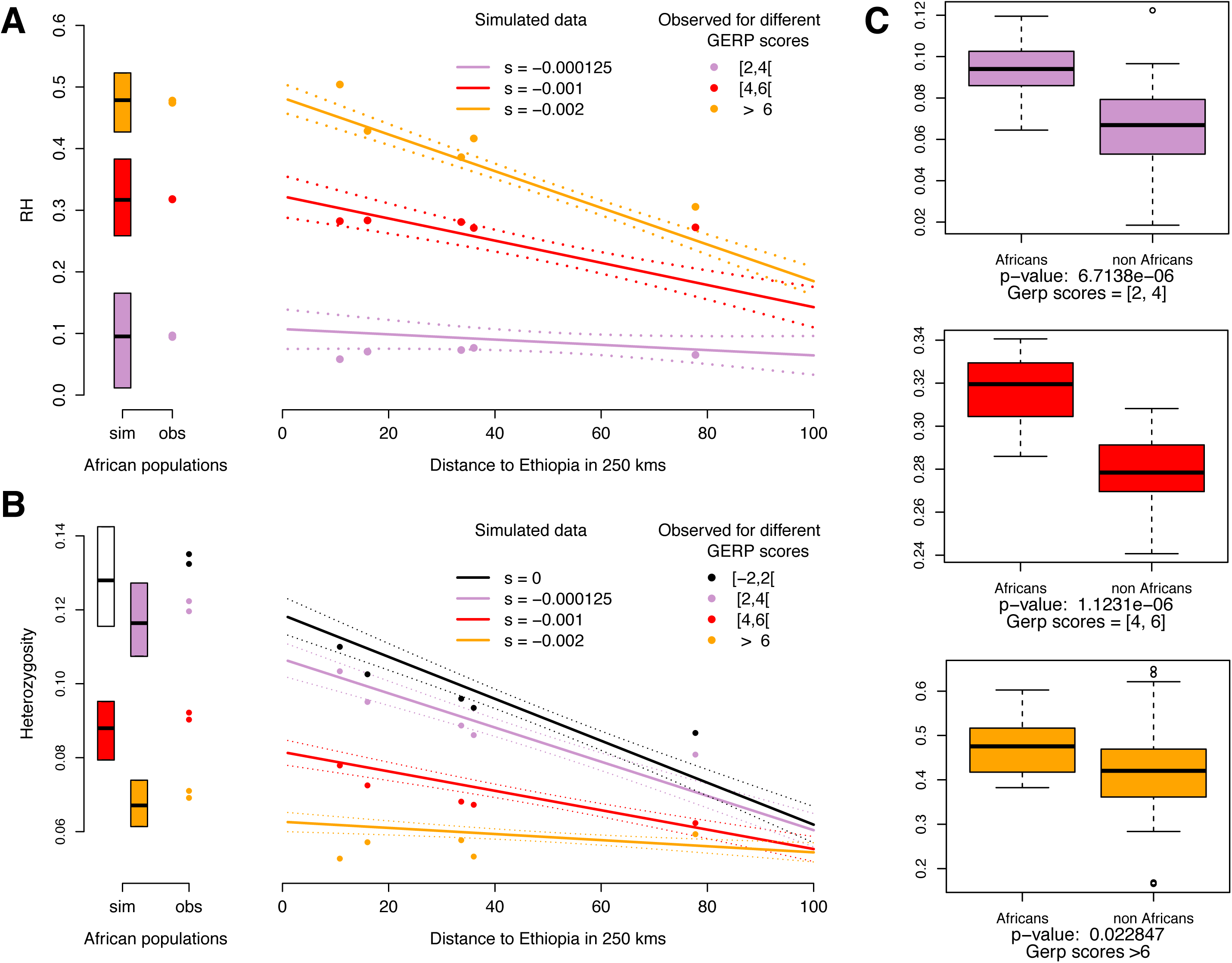
Heterozygosity under Range Expansion Simulations with Different Selection Coefficients. **A**) Observed and simulated patterns of the reduction of heterozygosity (*RH*). Selection coefficients used in the simulations are s= 0 (black), s= −0.000125 (lavender), s= − 0.001 (red), and s = −0.002 (orange). **B**) Colored circles show average expected heterozygosity for populations with ancestry from the OOA bottleneck. Solid lines show the regression lines obtained from simulations and dashed lines indicate 95% confidence intervals for the regression. The boxplots and colored circles on the left show the simulated heterozygosities in ancestral (i.e. African) populations, and the observed heterozygosity in our African dataset (San / Mbuti), respectively. **C**) Comparison of the distribution of RH between African and non-African individuals for different GERP categories, tested with a two-tailed Student t-test (see also **Fig. S15**).

*Evolutionary forces acting on heterozygosity*: In order to better understand which evolutionary forces have acted in different populations to shape their levels of genetic diversity, we define a new statistic “*RH*”. *RH* measures the reduction in heterozygosity at conserved sites relative to neutral heterozygosity, *RH* = (*H_neu_* - *H_del_*) / *H_neu_* where *H_neu_* indicates heterozygosity at neutral sites and *H_del_* at GERP score categories >2. *RH* can be seen as a way to quantify changes of functional diversity across populations relative to neutral expectations. For instance, a constant *RH* value across populations would suggests that average functional diversity is determined by the same evolutionary force(s) as neutral diversity, i.e. genetic drift and migration. In contrast, if *RH* changes across populations, it suggests that different evolutionary forces have shaped neutral and functional diversity, i.e. selection has changed functional allele frequencies.

In our dataset, *RH* is significantly larger in sub-Saharan Africans than in OOA populations across all functional GERP categories (**Fig. 4C**), indicating that selection has acted differently relative to drift between the two groups. The correlation between *RH* value and predicted mutation effect observed in Africa (**Fig. 4A**) confirms that purifying selection has kept strongly deleterious alleles at lower frequencies than in OOA populations. We then asked if there were significant differences across OOA population, as oriented by their distance from eastern Africa. Interestingly, we see that the OOA *RH* values do not depend on their distance from Africa for predicted moderate effect alleles (*p*=0.82, **Fig. S15**), suggesting that the frequencies of moderate mutations have evolved mainly according to neutral demographic processes during the range expansion out of Africa. In contrast, for strongly deleterious variants (“large” and “extreme” GERP categories) we see a significant cline in RH (*p*=0.01 and *p*=1.12×10–6, respectively, **Fig. S15**), which implies that purifying selection has also contributed to their evolution relative to demographic processes.

*Models of Dominance:* We next considered whether there is empirical evidence for non-additive effects for deleterious variants. Prior studies generally calculated “mutation load” by assuming an additive model, summing the number of deleterious alleles per individual, without factoring in whether a SNP occurs in a homozygous or heterozygous state. Determining an individual’s mutation load is, however, highly dependent on the underlying model of dominance (36) (see below for a formal definition of mutation load). For humans, Mendelian diseases tend to be overrepresented in endogamous populations or consanguineous pairings indicating that many of these mutations are recessive (37); Gao et al. (38) estimate 0.58 lethal *recessive* mutations per diploid genome in the Hutterite population. Gene conversion can also leads to differential burden of derived, recessive diseases alleles among populations (39). Even height, a largely quantitative trait appears to be affected by the architecture of recessive homozygous alleles in different populations (40).

To further clarify the impact of dominance, we compared the distribution of deleterious variants across genes associated with dominant or recessive disease as reported in Online Mendelian Inheritance in Man (OMIM) (41). We expect to see a lower proportion of large and extreme effect variants in genes with dominant OMIM mutation annotations, compared to genes with recessive OMIM mutation annotations. We tested this hypothesis with the HGDP as well as the much larger 1000 Genomes Phase 1 dataset (see *Supplementary Note,* **Fig. S18B**). We averaged the proportion of variants within each effect category and performed a Wilcoxon test to determine if the distribution of the proportion of large effect variants was different between dominant and recessive genes. In the HGDP dataset, we observed *p*= 0.06, and for the larger 1000 Genomes dataset, *p*= 0.03. Our results indeed show a significantly higher proportion of large effect variants in genes with recessive annotations, compared to genes with dominant annotations, suggesting that deleterious variants in the genome may tend to be recessive. However, we caution that in OMIM genes are annotated as dominant or recessive while dominance is a property of specific mutations, and therefore all deleterious variants in a gene will not necessarily have the same dominance coefficient. Nonetheless, our results are consistent with an interpretation that genes may have certain properties, e.g. negative selection against dominant mutations in crucial housekeeping or developmental genes, which influence the tolerable distribution of dominance among variants. We consider the effect of dominance (summarized by *h* which measures the effect of selected mutations in heterozygotes relative to homozygotes) on mutation load in the HGDP population samples given the observed differences in heterozygosity.

### Modeling the burden of deleterious alleles

We modeled three different scenarios to estimate the burden of deleterious alleles across populations The relationship between fitness *W* and load for a given locus *v* is classically defined (36) as:

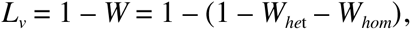

W_het_ = *gAa* × (1-*hs*) and W_hom_ = *gaa* × (1-*s*), where g*_Aa_* and g*aa* are the *observed* genotype frequencies of the heterozygotes and derived homozygotes respectively. The estimated population load (ignoring epistasis) is the sum of the load for all variants: *L_T_* = Σ*_v_ L_v_*. For each variant we assigned the selection coefficient inferred by the range expansion simulations according to its GERP score (see also Henn et al. (5)). Given that we do not know the distribution of dominance effects in human variation, we started by estimating the bounds for the mutation load for each population by considering two extreme scenarios: completely recessive and complete additive models for deleterious variants. We calculated *L_T_* for each HGDP population, **Fig. 5**. When all mutations are considered strictly additive, (*h*=0.5), values for mutation load are very similar across populations, with sub-Saharan African populations having the lowest mutational load (*L_T_* =2.83), followed by the Pathan and Mozabites, and finally the Asian and Native American populations showing the highest load (*L_T_* =2.89), **Fig. 5B**. We consider this model, as adopted in earlier studies, to demonstrate that even under an additive assumption, there is a statistically significant 1.7% difference in the spectrum of load between populations (*Supplementary Note,* **Fig. S24**). When all mutations are considered recessive (*h*=0), this model yields a much larger 45% difference in load (*L_T_* ranges between 1.27 and 1.85) between the San and the Maya, **Fig. 5A**. While this is surely an overestimate, it illustrates the broad range of potential values and consistent signal in the data for differences among population in estimated load. The mutation load under a recessive model is not explained by inbreeding, as measured by the cumulative amount of the genome in runs of homozygosity (cROH) greater than 1 Mb (r=0.27, p=0.55) (**Fig. S25**); this is because the African hunter-gatherers have relatively high cROH compared to other global populations, as is commonly observed in small endogamous populations (21, 42).

**Figure 5:**
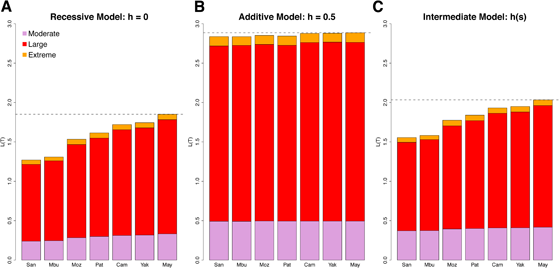
Estimates of Mutational Load in Seven Populations As a Function of Dominance assumptions. Total mutation load was summed over all annotated mutations in the exome dataset for the observed heterozygote and derived homozygote genotype frequencies in each population. The cumulative mutational load is shown in increased order from neutral to extremely deleterious mutations. Strongly deleterious mutations contribute the most to mutational load. Mutations were assigned an *s*, selection coefficient, based on their GERP score. A) *h*=0, recessive model; B) *h*=0.05, additive model; C) *h*(*s*), intermediate dominance model. For each selection coefficient, an *h* dominance coefficient was assigned based on the inverse relationship between *s* and *h*.

For the third scenario we used a model based on studies of dominance in yeast and *Drosophila* (19, 43, 44), in which there is an inverse relationship between selection and dominance (highly deleterious mutations tend to be recessive), and where *h* is sampled from a distribution following Agrawal and Whitlock (19). The maximal difference in load under this model was 30.8% (**Fig. 5C**), again between the San and Maya, and the minimum difference in load was 1%, between the Cambodians and Yakut. We note that the difference in relative fitness [e^-L(T)^] is much less than the difference in mutation load (i.e. a relative reduction of 79% in the San versus 87% in the Maya translates to a 8% difference between the two populations under the *h(s)* model, see also *Discussion*). As in the other modeled dominance scenarios, the majority of calculated mutational load is contributed by the large effect mutational category, as this category has a relatively strong selection coefficient and thousands of mutations (>4,000 on average per individual). Thus, this category contributes proportionally more to the total load, even though the extreme effect mutations have a higher selection coefficient. We note however that our assumed selection coefficients, particularly for the extreme effect, are somewhat lower than those obtained by other distribution of fitness effect (DFE) studies (16, 45) and simulations under an additive model results in even smaller selection coefficients (*see above*). As selection coefficients are the same across populations in our calculations, *s* will affect the absolute value of load but not relative differences across populations.

## Discussion

Two primary demographic signals are reflected in human genetic data from non-African populations First, a major five-to-tenfold population bottleneck is associated with the Out-of-Africa dispersal(s) (46–48). Second, the distribution of genetic diversity among non-African populations is characterized by a decrease in heterozygosity proportional to geographic distance from northeastern Africa. A model of serial founder effects in the ancestral populations of Eurasia, Oceania and the Americas has been posited as the most likely model for explaining the systematic variation in genetic diversity across this geographic range for humans (25, 26), as well as commensal human species (49, 50). By directly ascertaining genomic variation in over 50 individuals from 7 populations, we observe a clear cline of genetic diversity as a function of distance from Africa supporting evidence for a serial founder effect model. We also observe differences in the amount of predicted deleterious variation across populations. These differences appear to result from the genetic drift of existing deleterious variants to higher frequencies during the sequential range expansion after the Out-of-Africa exit (**Fig. 3B**). Clines in heterozygosity for the different mutational effect categories can be reproduced by spatially explicit simulations with negative selection and recessive mutations (**Fig. 4**, see also codominant simulations, **Fig. S16**). Although both moderate and large effect deleterious mutations have evolved under negative selection in Africa (**Fig. 4C, Fig. S15**), many predicted moderate variants have evolved as if they were neutral in non-African populations. However, selection has remained a major force during the OOA expansion for strongly deleterious variants.

***Impact of the Out-of-Africa Bottleneck***: There is an ongoing debate on whether selection has been equally or more efficient in African vs. non-African populations due to the major bottleneck that occurred in the ancestors of OOA populations (10, 12, 13, 35). Two studies found no significant differences in mutation load between European-American and African-Americans under an additive model with two classes of alleles: deleterious and neutral (12, 13, 33). Fu et al. (11) identified small but significant differences in the average number of alleles and the SFS, potentially due to a different algorithm for predicting mutation effect than earlier studies. We argue that estimates of the efficacy of selection should take into account not only the number of mutations per individual but the predicted severity of mutational effect. Here, we classify mutations into four categories and find differences across populations in some, but not all, mutational categories. For variants that have putatively moderate (2< GERP < 4) or extreme deleterious effect (GERP > 6), we do not see a significant difference between African and non-African populations in the number of mutations per individual. Significant per individual differences are only observed for the intermediate large effect category. We used PhyloP scores (51) as an alternative measure of conservation to verify our main results (see **Fig. S26**). We found qualitatively very similar patterns for both the spatial distribution of the number of derived homozygous sites per individual (**Fig. S26 A**) as well as the number of derived alleles per individual (**Fig. S26 B**), suggesting that our results are robust to the choice of prediction algorithm that is used to estimate deleteriousness of mutations.

We note that the observed differences between populations are relatively small compared to the within population variance (**Fig. 2**). Nonetheless, a novel measure of the efficacy of selection, *RH*, is significantly different across all three mutational categories (**Fig. 4C, Fig. S15**) between sub-Saharan Africans and non-Africans in our dataset. That is, the observed heterozygosity at deleterious loci is greater in non-Africans than Africans - after correcting for neutral genetic diversity in each group. This is particularly significant for moderate and large effect mutations, in agreement with theory that would suggest that differences in purifying selection will primarily emerge for variants at the *N_e_s* boundary.

***Serial Founder Effects / Range Expansion:*** Several simulation studies have attempted to characterize the distribution of deleterious alleles under Out-of-Africa demographic scenarios. Some simulations focused on differences in the cumulative number of deleterious alleles per individual; others focused on differences in the proportion of segregating alleles within a population that are deleterious. Lohmueller et al. (10) found that a long bottleneck lasting more than 7,500 generations (>150 *Ky*) could produce the excess proportion of deleterious mutations observed in European-Americans. A bottleneck model with subsequent explosive growth has also been proposed to explain the proportionally greater number of non-synonymous or deleterious mutations in Eurasian populations (52, 53). As a consequence, deleterious mutations accumulate in populations during the expansion process. Simons et al. (12) tested a long bottleneck and subsequent population expansion model, contrasting African and non-African populations and found no evidence that human demography played a role in the differential accumulation of deleterious alleles per individual.

A recent theoretical study of spatial range expansions (*i.e.* a model similar to geographic serial founder effects) showed that strong genetic drift at the wave front of expanding populations decreases the efficiency of selection (32). Under a spatial range expansion model, deleterious variants, unless they have a large selection coefficient, should evolve as if they are neutral on the wave front (32), and their overall frequency should therefore not change much during the range expansion (7). The loss of deleterious variants at some loci should be compensated by an increase of their frequencies at other loci. The frequency of deleterious homozygotes should therefore increase with distance from Africa, which is observed here in the rightward shift of the SFS in OOA populations (**Fig. 3**), except for the most evolutionarily constrained sites. We can address the question of whether this increased frequency is driven entirely by drift and gene surfing, or by differential selection in non-African populations by considering the spatial distribution of the *RH* statistic (**Fig. 4C**). The fact that *RH* does not change among OOA populations for moderately deleterious alleles suggests that they have evolved as if they were neutral alleles during the expansion and that selection has not yet purged the deleterious mutations that increased in frequency. In contrast, extremely deleterious alleles (GERP > 6) exhibit similar heterozygosity in all OOA populations, suggesting that they are subject to similar levels of purifying selection in these populations. The remaining deleterious alleles (4<GERP<6) present an intermediate pattern, implying that both drift and selection have acted on this category of sites.

A recent controversy concerns whether there are differences in the efficacy of purifying selection between African and non-African populations (6, 12, 13). It is difficult to discuss our results in the context of this controversy because there is no generally accepted definition of “efficacy of selection”, and different definitions will lead to different interpretations (4). We therefore prefer to interpret our results in the context of our spatially explicit model of range expansions, and the relative roles of drift and selection in this model. Recurrent founder events should contribute to a decrease in the effective population sizes with distance from Africa, and it is commonly assumed that selection will become weaker with smaller effective population sizes. However, reducing the impact of a range expansion to a simple gradient in effective size, and thus to a decrease of the efficacy of selection can be misleading. Diversity-based estimates of *N_e_* are not necessarily informative about the strength of selection in non-equilibrium scenarios because estimates of *N_e_* may lag behind recent demographic changes (e.g., (54)). Rather, if one considers that deleterious alleles were kept at low frequencies by purifying selection in ancestral African populations, those that increased in frequency by gene surfing during the OOA expansion also became more accessible to subsequent selection, especially for those alleles that were recessive. The observed cline in RH for large effect mutations is more compatible with an unequal purging of deleterious variants by selection. Indeed, selection will have had less time to act on newly formed populations that are further away from Africa, and it will also operate more slowly on populations that have less diversity and therefore lower inter-individual differences in fitness. Furthermore, the fact that our simulations can reproduce the observed pattern with spatially uniform population sizes and strength of selection against deleterious mutations implies that the simulated gradients in *RH* in **Figure 4A**, as well as the increased number of deleterious homozygous sites, is not the consequence of reduced strength of selection away from Africa. Rather, it is caused by increased drift during the expansion, as well as by differential purging of deleterious mutations after the expansion.

***The importance of dominance:*** Multiple modeling assumptions are crucial when considering the burden of deleterious alleles across populations. In addition to the selection coefficients, the assumed dominance terms are critical. An estimated 16% of Mendelian diseases are known to be autosomal recessive (estimated from the OMIM) and many contribute significantly to infant mortality. Due to the difficulty of detecting recessive diseases, unless they are extremely damaging, there are potentially many more disease mutations that have an *h* coefficient less than 0.5. Autosomal recessive diseases appear to be more frequent than autosomal dominant diseases (55) and even mildly deleterious mutations are predicted to have a mean *h* of 0.25 (56). Although formal calculations of genetic load require multiple assumptions, we demonstrate that differences in calculated load across human populations are primarily sensitive to assumptions about dominance, as expected given the increased extent of homozygosity in OOA populations.

We have modeled deleterious mutations as having variable *h* coefficients. While strongly deleterious mutations are likely recessive, dominance for weakly deleterious mutations is particularly problematic to estimate because there is less power to measure weak effects and *h* may be upwardly biased in model organism competition experiments (19). When sampling *h* coefficients under our model, we allowed weakly deleterious mutations to be assigned a coefficient *h*>0.5, but this had little effect on mutational load because the bulk of the load was contributed by large effect variants. However, a fraction of strongly deleterious mutations are clearly dominant as ascertained from disease studies and future work may need to model different mixtures distributions on *h*. We also note that the absolute mutational load is two-fold higher under an additive model than under a recessive model (**Fig. 5**), as expected from theory (36).

***Estimates of Mutational Load:*** We estimate that there are differences in mutational burden calculated using a formal load model, among extant human populations, particularly if we depart from a simple additive assumption. We found that the change in mutation load between sub-Saharan African populations versus Native American populations (the two ends of the range) were significantly different at p<0.05 under recessive, partially recessive and additive models (**Fig. S24**). Mutational load under a fully or partially *recessive* model is ten to thirty percent greater in non-African populations (**Fig. 5A**), as the result of higher homozygosity from the legacy of the OOA bottleneck across all (deleterious) mutation categories [e.g. L_T(Mbuti)_ = 1.59, L_T(Yakut)_ = 1.95 under the *h*(*s*) model]. All populations carry significant load, relative to a population with the alternate, ancestral allele genotype. Under a model where fitness differences are determined *only* by genotype and environments are equal across individuals, the relative fitness [e^-L(T)^] of 0.204 for the Mbuti indicates a reduction in fitness of 79.6%, whereas a relative fitness of 0.142 for the Yakut indicates an 85.7% reduction. These fitness differences are relatively small, even under a partially recessive model.

Although illustrative, such models of load have important limitations. The mutations identified in this dataset have not been functionally characterized and are predicted to be deleterious based on degree of sequence conservation. The assumed selective coefficients across GERP categories are fit based on a recessive model, which is not applicable to all sites. However, while different selection coefficients will change the values of load in our calculation, it will not change the relative difference among populations because the same set of coefficients were applied to all populations (5). If mutations have different fitness effects across heterogeneous global environments, then the values of mutation load will change. Indeed, a proportion of the alleles may be locally adaptive, or neutral, and hence the sign of the selection coefficient for the mutation would be misestimated in our analysis. For example, the Duffy null allele is classified as a large effect mutation using GERP (RS=4.27) and is found at high frequency in western Africa; however, it has likely increased in frequency due to positive selection as a response to malaria (57). Recent genome-wide studies have stressed the paucity of selective sweeps in the human genome (35, 58, 59); only 0.5% of non-synonymous mutations in 1000 Genomes Pilot Project were identified has having undergone positive selection. Others have emphasized evidence for pervasive adaptive selection (60, 61) and a variety of studies have identified specific beneficial alleles locally adapted to high altitude, immune response and pigmentation (62–64). We considered local adaptive evolution by examining highly differentiated alleles in our dataset, i.e. alleles that differ by 80% in frequency between a pair of populations, indicative of a strong local adaptation. We find that highly differentiated alleles have the same GERP score distribution as non-differentiated alleles, indicating there is little reason to believe that most large and extreme effect mutations have been subjected to strong local adaptation (**Fig. S20**, also (65)). We conclude that the raw, calculated mutational burden may differ across human populations, although the effects of positive selection, varying environments and epistasis have yet to be explored, and remain a significant challenge to fully understanding mutational burden.

## Conclusions

A major difference between our work and previous results is the interpretative framework we present, which underlines the role of range expansions out of Africa to explain patterns of neutral and functional diversity Whereas previous comparisons between African and non-African diversity attributed the observed increased proportion of deleterious variants in non-Africans to the Out of Africa bottleneck (10), our study shows that a single bottleneck is not sufficient to reproduce the gradient we observe in the number of deleterious alleles per individual with distance from Africa (**Fig. 2**). Taking into account the range expansion of modern humans (66) sheds new light on this apparent controversy. Finally, we note that recent simulation work (4) suggests that the impact of a bottleneck on the efficacy of natural selection depends critically on the distribution of fitness and dominance effects as well as post-bottleneck demographic history. While these models and parameter choices clearly affect the interpretation of the pattern of deleterious alleles across populations, we find empirical evidence for significant differences in deleterious alleles as tabulated by a variety of statistics across the spectrum of human genetic diversity.

## Materials and Methods

### Samples and Data

Aliquots of DNA isolated from cultured lymphoblastoid cell lines were obtained from Centre d’Étude du Polymorphisme and prepared for both full genome sequencing on Illumina HiSeq technology and exome capture with an Agilent SureSelect 44Mb array 101 bp read-pairs were mapped onto the human genome reference (GRCh37) using a mapping and variant calling pipeline designed to effectively manage massive amounts of short read data. This pipeline followed many of the best-practices developed by the 1000 Genomes Project Consortium (34).

### Variant Annotation

Ancestral state was inferred based on orthologous regions in a great ape and rhesus macaque phylogeny as reported by Ensembl Compara and used by the 1000 Genomes Project. To determine the biological impact of a variant we used GERP score (30) a measure of conservation across a phylogeny. Positive scores reflect a site showing high degree of conservation, based on the inferred number of “rejected substitutions” across the phylogeny. GERP scores were obtained from the UCSC genome browser (http://hgdownload.cse.ucsc.edu/gbdb/hg19/bbi/All_hg19_RS.bw) based on an alignment of 35 mammals to human. The allele represented in the human hg19 sequence was not included in the calculation of GERP RS scores. The human reference sequence was excluded from the alignment for the calculation of both the neutral rate and site specific ‘observed’ rate for the RS score to prevent any bias in the estimates. In addition to GERP, we also used PhyloP scores (51) as measures of genomic constraint during the evolution of mammals. We used the PhyloP_NH_ scores computed in Fu et al. (11) from the 36 eutherian-mammal EPO alignments (available in Ensembl release 70 (67)), which is also computed without using the human reference sequence.

### Classification of mutation effects by GERP scores

Variants were classified as being “neutral”, “moderate”, “large” or “extreme” for GERP scores with ranges [-2,2], [2,4], [4,6] and [6,max], respectively. The use of four “bins” of GERP scores simplifies the range expansion simulations performed for distinct selection coefficients. For every individual the total number of derived deleterious counts found in homozygosity (i.e. 2 x HOM), and the total number of deleterious counts (i.e. HET + (2 × HOM)) within each category was recorded.

### Individual-based simulations

To simulate changes in heterozygosity, we modeled human range expansion across an array of 10x100 demes (32). After reaching migration-selection-drift equilibrium, populations expand into the empty territory, which is separated from the ancestral population by a geographical barrier, through a spatial bottleneck (**Fig. S21**). After 3,000 generations, we computed the average expected heterozygosity for all populations. The migration rate and selection coefficients were adjusted to generate heterozygosity consistent with the observed data, without formally maximizing the fit.

### Calculating Load

Mutational load was calculated following Kimura (36), but using observed genotype frequencies instead of inferring them from Hardy-Weinberg based on the allele frequencies. In this way, the fitness of the heterozygotes and the homozygotes will be: W_het_ = Aa × (1-*hs*) and W_hom_ = aa × (1s), where *Aa* and *aa* are the genotype frequencies of the heterozygotes and derived homozygotes respectively. The fitness for a given variant will be relative to that of the ancestral variant, which for numerical convenience is set to 1. The relationship between fitness and load is: *L_v_* = 1 − *W* = 1 − (1 − *W_he_*_t_ − *W_hom_*), and the total population load is the sum of the load for all variants, *L_T_* = Σ*_v_ L_v_*.

## Author Contributions

C.D.B, L.E., B.M.H. and J.M.K. conceived of and designed the experiments. B.M.H., L.R.B., A.R.M., B.K., S.M., M.L., S.P., I.D. and J.M.K. performed analyses. H.C. and M.S. contributed materials. B.M.H., L.R.B., J.M.K, S.P., C.D.B, and L.E. wrote the manuscript.

## Acknowledgements

We thank Chris Tyler-Smith, David Reich, Yuval Simons, Spencer Koury and Simon Gravel for helpful discussion. L.R.B was supported by a Beatriu de Pinós Programme Fellowship. C.D.B. and B.M.H acknowledge support from NIH grant 3R01HG003229; J.M.K. was supported by NIH grant DP5OD009154. S.P. and I.D. were supported by a Swiss SNSF grant 31003A-143393 to L.E.

**Data deposition**: Data have been deposited in the SRA [SRP036155] and in dbSNP. VCF files are freely available on the Henn Lab website: http://ecoevo.stonybrook.edu/hennlab/data/.

